# Development of a Polyelectrolyte Complex Scaffold and its specific cell seeding method as a tool for liquid cancers drug screening

**DOI:** 10.64898/2026.04.30.722037

**Authors:** S. Grossemy, S. Cadot, M. Farno, S. Cavalie, B. Sallerin, L. Ysebaert, A. Quillet-Mary, Fullana S. Girod

## Abstract

This study focuses on the development of 3D culture model dedicated to liquid cancers drug screening. The challenge addressed was to effectively retain non adherent small cells within a 3D-scaffold with tailorable mechanical properties, while proposing a fast and effective tool for drug screening. To that aim, we developed a macroporous alginate–chitosan polyelectrolyte complex (PEC) scaffold combined with a low-viscosity alginate (LVA) cell seeding solution. We hypothesized that LVA could undergo in situ pore gelation via calcium ions retained from the PEC fabrication process, enabling effective retention and homogeneous cell distribution, leading to an improved platform for drug screening and personalized medicine.

First, we evaluated scaffold suitability for LVA infiltration and gelation. Microtomography revealed a highly porous architecture (98%) enabling LVA homogeneous penetration and complete gelation within 30 min, as confirmed by SEM, microscopy, rheology, and micro-rheology.

Next, we assessed cell retention and biocompatibility using primary human chronic lymphocytic leukemia (CLL) cells. LVA-assisted seeding increased cell density 2.6-fold compared to medium alone, with homogeneous distribution, >80% viability over 7 days, and preserved differentiation into nurse-like cells.

Finally, we demonstrated a proof of concept for drug screening. The Alginate-PEC scaffold (A-PEC scaffold) supported both qualitative live/dead imaging and rapid quantitative viability measurement with the Alamar Blue assay. Drug responses reproduced microenvironment-dependent protection effects observed in vivo.

This integrated scaffold and seeding method provides a promising 3D platform for in vitro liquid cancer studies and drug screening on patient-derived hematological cancer cells.

**Graphical abstract:** 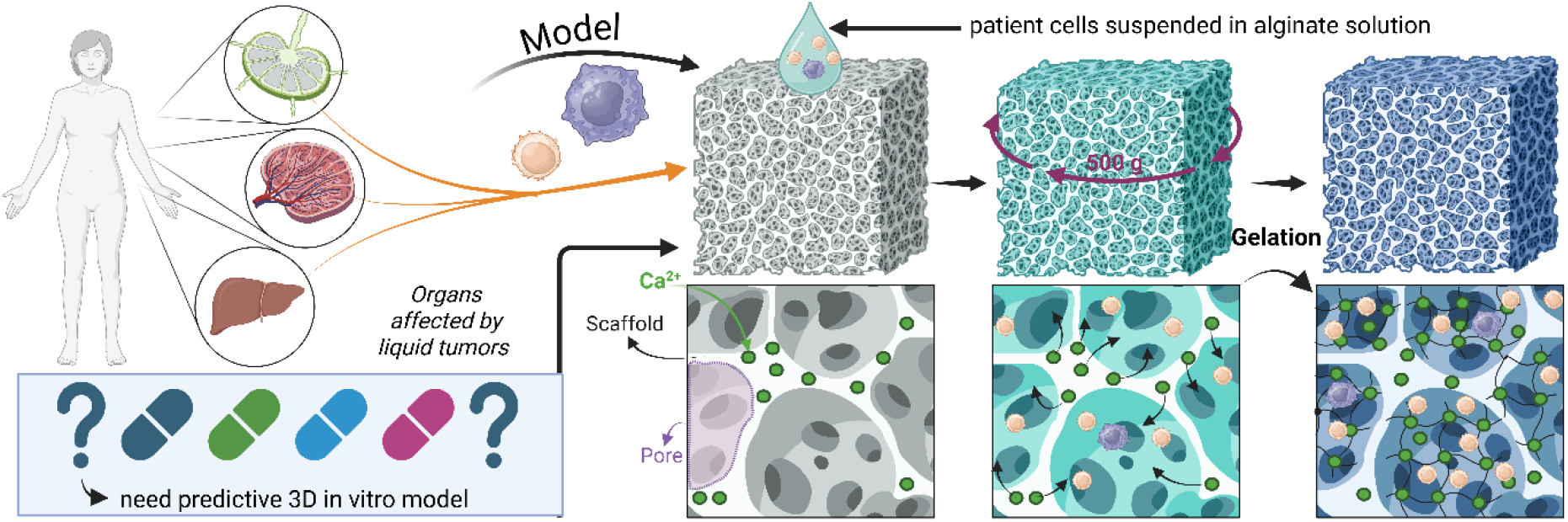

## 2 Introduction

Liquid tumors, such as leukemia, present significant challenges in treatment due to the variability in patient responses to therapy [1, 2]. With a wide range of available treatments, including chemotherapy, immunotherapy, and targeted therapies, it becomes crucial to develop *in vitro* models that can more accurately predict patient-specific responses. These models are essential for advancing predictive medicine, where the effectiveness of therapeutic strategies can be assessed in a personalized manner using patient-derived cells. Currently, the traditional two dimensional (2D) *in vitro* model is the most common strategy for cancer therapeutics screening. Although relatively cheap and fast, current 2D models struggle to replicate complex mechanical and biochemical conditions, such as stiffness or tumor microenvironment, limiting their utility in guiding treatment as a personalized drug screening tool. Thus, the development and validation of 3D methods to overcome these limitations is a field of active expansion. Different scaffold-free and scaffold-based approaches have been developed to better integrate tumoral microenvironment complexity in 3D, such as spheroids, organoids, hydrogels, decellularized and material-based scaffolds [3-6].

The application of 3D approaches in hematology is particularly challenging. Leukemia, such as Chronic Lymphocytic Leukemia (CLL), is characterized by the accumulation of small quiescent tumoral lymphocytes in the blood, and proliferative leukemic cells in secondary lymphoid organs (SLO), where leukemic cells survival is highly dependent on cross-talk with cells from the microenvironment (T lymphocytes, Nurse Like cells (NLC)) and on the stiffness of the organ [7, 8], often limiting therapeutic drug efficacy *in situ* [9-11].

These characteristics present a dual challenge for the development of *in vitro* 3D models. The first challenge is the retention of non-adherent circulating cells within a 3D system, which typically requires encapsulation within a hydrogel [5, 12, 13]. The second challenge arises from the diversity of tissue stiffnesses affected by the disease, necessitating a model that can accurately reproduce these mechanical features. Therefore, the real challenge lies in the smart combination of cell encapsulation/seeding techniques that ensure cell retention within a hydrogel while simultaneously achieving mechanical properties that reflect those of the affected tissues.

Unfortunately, conventional hydrogels used for cell encapsulation often lack the mechanical strength required to mimic the stiffness of the tumor microenvironment. This is the case for alginate and chitosan hydrogels, although their excellent biocompatibility attributed to their biomimetic resemblance to the extracellular matrix [14]. Interestingly, they can be combined as polyelectrolyte complexes (PEC) which have shown strong potential for various biomedical applications, depending on their architecture [15-17].

Recently, we developed a platform of chitosan-alginate PEC scaffolds and demonstrated their biocompatibility [18]. By adjusting the alginate/ chitosan ratio, a synergistic enhancement of their respective mechanical properties can be achieved, thus offering a variety of mechanical properties fitting with the mechanical properties of soft tissues such as lymph nodes [15, 17, 19]. A combination of foaming and drying conditions allows modulation of 3D architecture, particularly pore size and distribution, enabling in-depth cell seeding and tailorable cells confinement. However, such scaffolds fail to retain non adherent cells, which could be a serious limitation when studying circulating blood cells.

Based on these considerations, we hypothesized that combining a macroporous alginate–chitosan PEC scaffold with a low-viscosity alginate (LVA) cell seeding solution could address both challenges. More specifically, we propose that calcium ions intrinsically retained within the PEC scaffold could trigger *in situ* gelation of the LVA within the pores, enabling efficient cell retention and homogeneous distribution, while preserving the mechanical properties of the scaffold.

To explore this hypothesis, we first investigated the ability of the LVA seeding solution to infiltrate the porous network and undergo *in situ* gelation. We then assessed cell retention and viability using primary chronic lymphocytic leukemia (CLL) cells, as well as their capacity to differentiate into nurse-like cells. Finally, we evaluated the relevance of this platform for drug screening by comparing therapeutic responses with conventional 2D cultures and 3D spheroids.

## 3. Materials and Methods

All chemicals were purchased from Sigma Aldrich (St. Louis, MO) unless otherwise specified. Chitosan medium viscosity, sodium alginate low and medium viscosity (from brown algae) powders were used without any further modification. Ibrutinib and venetoclax were purchased from Selleckchem.

### 3.1. Preparation of Polyelectrolyte Complex (PEC) Scaffolds

PEC scaffold synthesis: a 4.5 % chitosan in 0.25M acetic acid solution and a 3 % medium viscosity alginate solution were prepared separately by mixing for 30 min at 1000 rpm (RZ-2041, Heildolph, Germany). The resulting solutions were then mixed together to obtain a 40/60 alginate/chitosan w/w ratio in presence of 0.9% w/w sodium bicarbonate and 1% w/w Montanox 20 at 1600 rpm for 30 min. The resulting foam was cast in 96-well cell culture plates and frozen at -20°C for 12 h. The samples were then lyophilized, crosslinked in 0.1 M CaCl_2_ and 0.25 M acetic acid for 1 hour at room temperature, washed with deionized water with increasing ethanol concentration up to 100%, and then critical point dried (E3000 Series Critical Point Dryer, Quorum Technologies).

### 3.2. X-ray tomography of the scaffold

The X-ray microtomography acquisitions were performed using a Nanotom 180 system manufactured by Phoenix/GE, located at the FERMat Research Federation of CNRS FR 3089 in Toulouse. A diamond target was utilized for the X-ray source. The settings for both analyses were set to a voltage of 50 kV and a current of 350 μA. During the acquisition process, 2000 steps were taken over a 360-degree rotation of the sample. For each angular step, the first image was discarded to eliminate any potential issues with remanence. Subsequently, four images, each with an exposure time of 750 ms, were acquired and averaged. For the rat tendon sample, the source-to-object distance was set to 7.2 mm, and the source-to-detector distance was 250 mm, resulting in a voxel size of 1.5 μm. For the bovine tendon sample (larger in size), the source-to-object distance was set to 16 mm, and the source-to-detector distance was 300 mm, resulting in a voxel size of 2.8 μm.

### 3.3. Low-viscosity alginate (LVA) solution preparation and sterilization

Low-viscosity alginate (SLBX8356 Sigma) solution was prepared at a 0.5% (w/v) concentration by dissolving it in a NaCl solution (0.9% in water) under sterile conditions. The solution was mixed using a motorized paddle for 60 minutes to achieve homogeneity (1000 rpm). After preparation, the alginate solution underwent sterilization using either filtration or autoclaving. For filtration, the solution was passed through a 0.22-micrometer sterile filter to remove potential contaminants. Both sterilization methods were evaluated to confirm the preservation of the alginate’s integrity and functionality for subsequent applications.

### 3.4. Rheological Properties of low Mw alginate Solutions

Rheological measures were performed to assess the flow properties of both sterilized and non-sterilized alginate solutions using a rotational rheometer equipped with a parallel plate geometry. Alginate solutions were prepared at a concentration of 0.5% and equilibrated at room temperature before testing. The gap between the plates was set to 52 µm, and a frequency sweep was conducted at a constant strain of 1% to determine the shear modulus over frequency range. Shear stress and shear strain data were collected to generate shear stress versus shear strain rate curves, while viscosity as a function of shear strain rate was also analyzed across the same frequency range. All measurements were carried out at a controlled temperature of 37°C to ensure consistent and accurate evaluation of the rheological behavior.

### 3.5. Low-viscosity alginate FITC functionalization and imaging

A total of 300 mg of low viscosity alginate was dispersed in 100 mL of MES buffer (2-morpholinoethanesulfonic acid, 50 mM MES, pH 6) containing 10% ethanol. Following this, an EDC (1-ethyl-3-(3-dimethylaminopropyl)carbodiimide hydrochloride) and NHS (N-hydroxysuccinimide) solution was added, maintaining the molar ratios of [EDC]/[polymer repeat units] = 1 and [NHS]/[EDC] = 1 (1.5 10^-5^ mol/L) the solution was stirred for 15 min at room temperature. Subsequently, a solution of FITC (fluorescein isothiocyanate, Interchim UP017396) in ethanol was introduced to label 1/1000th of the alginate carboxyl groups (1.042 mL were added at a concentration of 5mg/mL in FITC). The reaction mixture was stirred overnight, protected from light.

The resulting FITC-labelled polymer solutions were dialyzed sequentially: first against 0.05 M sodium bicarbonate, then against 0.05 M sodium chloride for 72 hours, and finally dialysis step was repeated with NaCl solution twice for 24 hours. The pH of the Alg-FITC solution was adjusted to 7.4.

The solution of functionalized low-viscosity alginate was prepared at a concentration of 0.5%, as described in Section 1.1. A total of 10 µL of this solution was carefully loaded onto the precasted PEC in a 96-well plate. The plate was then centrifuged at 400 RPM for 4 minutes to ensure proper distribution of the alginate over the PEC surface. For control purposes, a PEC without any alginate was used to account for the background fluorescence of the PEC itself.

### 3.6. Cryo-MEB

PEC samples were prepared by casting in a 96-well plate. A low-viscosity alginate solution (10 µL) was applied on top of the PEC and centrifuged at 400 × g for 1 minute to incorporate the alginate into the PEC structure.

After centrifugation, the PEC samples were snap-frozen by directly plunging them into liquid nitrogen for 10 min. This rapid freezing was crucial to maintain the structural integrity of the samples. Once frozen, the PEC samples were manually fractured to expose internal structures. The fractured samples were then mounted onto cryo-compatible metal stubs using conductive carbon paste to ensure proper adhesion during imaging. The mounted samples were transferred to the Cryo-SEM stage for imaging. No additional sputter coating was applied. Imaging was performed under cryogenic conditions to capture the structure of the alginate-incorporated PEC samples.

### 3.7. Monitoring of low Mw alginate In Situ Gelation by Dynamic microrheology measurement

Microrheology was employed to monitor the micro-dynamics of the alginate gelation process, utilizing a RMaster (Formulaction, Toulouse, France). The PEC were mold in 24 well plate, and a 0.5% LVA solution was subsequently applied onto the PEC matrix (10 ul). To enhance the incorporation of alginate into the PEC structure, centrifugation was performed following the application (1 min 400g). Immediately afterward, the measurement container was placed into the RMaster to assess molecular agitation and track the gelation process in real time. This allowed for the continuous monitoring of changes in the sample’s optical properties during gel formation, providing valuable insights into the dynamics of gelation.

### 3.8. Monitoring of low Mw alginate In Situ Gelation by Rheology Measurement

Rheological measurements were carried out using a stress-controlled rheometer (Rheostress RS75, HAAKE, Germany) with a parallel-plate geometry (plate diameter:20 mm) on fully rehydrated specimens in a thermostated immersion cell. For sample preparation, the PEC was molded into 6 wells plates and subsequently cut using scissors to fit the exact dimensions of the parallel-plate geometry (20 mm). Once prepared, a solution of sterile-filtered alginate (0.5% w/v) was applied onto the surface of the PEC. The sample was then centrifuged at 400 G for 1 minute to ensure proper integration of the alginate into the PEC structure. Following centrifugation, the sample was immediately placed under the rheometer for measurement. For the control the same process was followed using RPMI solution. The applied volume of alginate or RPMI applied on the PEC previous to centrifugation was 2 mL.

The gap between the plates of the rheometer was set to a value of 4 mm. A time sweep was performed to monitor the increase in the storage modulus (G’) of the PEC-alginate system, indicating the progression of gelation. The amplitude of the oscillation was set to 0.1%, with a frequency of 10 Hz, to capture the rheological properties throughout the gelation process. Measured were conduct in immersion in RPMI

### 3.9. Cell Seeding and Culture

Peripheral blood samples from untreated Chronic Lymphocytic Leukemia (CLL) patients were obtained from the Hematology Department with informed consent and referenced in INSERM cell bank. According to French law, INSERM cell bank has been registered with the Ministry of Higher Education and Research (DC20131903) after being approved by an ethic committee. Clinical and Biological annotations of the samples have been reported to the Comité National Informatique et Liberté.

Fresh peripheral blood mononuclear cells (PBMC) were isolated by density gradient sedimentation (Ficoll-Hypaque, GE Healthcare), washed and resuspended at 200 x 10^6^ cells/mL in RPMI-1640 10%FCS (2D and 3D experiments) or in 0.5% LMW Alginate (A-PEC scaffolds experiments).

#### 2D experiments

10µL of cells were seeded in 96-well flat-bottom plates. 170 µL of RPMI-1640 10% FCS were added in each well.

#### 3D experiments

10µL of cells were seeded in 96-well Nunclon Sphera round bottom plates (ThermoFisher scientific). 170µL of RPMI-1640 10%FCS were added in each well. Plates were centrifuged 1 min at 400g to ensure the formation of spheroids.

#### A-PEC scaffolds experiments

Scaffolds were placed in 96-well flat-bottom plates. 10 µL of cells (in 0.5% LMW Alginate, or in medium, depending of the experiment) were seeded at the top of the scaffold. Plates were centrifuged 1 min at 400g without brake, to ensure the penetration of the cells. After 20 min, 170 µL of RPMI-1640 10%FCS were added in each well.

For all experiments (each condition in triplicate), cells were left at 37°C 5% CO_2_ for one night before adding or not drug treatments. For long-term experiments, cells were seeded as described above and left 7 days at 37°C 5% CO_2_ to ensure Nurse Like Cells (NLC) differentiation [11, 20] before drug treatment during 7 days more. Controls without NLCs were experiments at 7 days culture.

### 3.10. Live/Dead Cell Staining and Confocal Imaging

A staining solution was prepared by dissolving ethidium homodimer-1 (ETH) at a concentration of 4 µM and calcein AM (ThermoFisher) at 2 µM in 37 degrees DPBS ca2+. The scaffold was initially rinsed with 37 degrees DPBS ca2+ at 37°C for 10 minutes to remove any residual media and debris. After the rinsing step, the DPBS was removed, and the staining solution was applied to the scaffold, followed by incubation at 37°C for 45 minutes to ensure thorough staining. Prior to imaging, the scaffold was rinsed twice with DPBS containing calcium ions at 37°C to eliminate any unbound staining reagents. Confocal microscopy was then used to capture the fluorescent signals corresponding to live cells (green, calcein AM) and dead cells (red, ETH), allowing for quantitative analysis of cell viability and distribution within the scaffold.

Confocal laser scanning microscopy (Zeiss LSM 880, Carl Zeiss) was utilized to assess cell distribution and quantify cell populations within the PEC-alginate scaffolds. Z-stack acquisition was performed to a depth of 150 µm with a step size of 10 µm, capturing a total of 16 optical slices per sample. Each Z-stack was acquired using a 10× objective lens, combined with a 9 × 9 tile scan to ensure full coverage of the scaffold.

Following acquisition, the image stacks were processed using IMARIS software (Bitplane), which enabled 3D reconstruction of the scaffold and accurate cell counting. Automated spot detection in IMARIS was employed to quantify the total number of cells per scaffold, adjusting for cell size and signal intensity thresholds. Quantifications were performed in triplicate, and data were expressed as mean cell count ±standard deviation (SD).

### 3.11. Cell viability assessment in A-PEC scaffolds using the Alamar Blue assay

For drug screening, cells were seeded in 2D, 3D or A-PEC scaffold conditions (in triplicate). After one night at 37°C 5% CO_2_, ibrutinib or venetoclax (10µL) or complete medium (10 µL, positive control) were added to the cells. For short-term cultures (48hrs), ibrutinib was used at a final concentration of 1µM or 10µM and venetoclax at a final concentration of 1nM or 10nM; for long-term cultures (7 days), final concentration was 0.5µM for ibrutinib and 0.5nM for venetoclax [10, 21]. Plates were incubated at 37°C 5% CO_2_. Then, viability of cells was determined using Alamar Blue assay. Briefly 20 µL of Alamar Blue solution was added in each well. After an incubation of one night at 37°C5% CO_2_, 100 µL of supernatant (from each well) were transferred to a black-bottom 96-well plate to measure fluorescence values on a microplate reader following manufacturer instructions (Clariostar instrument). Percentage (%) of viable cells was calculated based on negative control (medium or alginate alone), and positive controls (cells without drugs). Statistical analysis of biological data was done using one-way Anova *p < 0.05; **p < 0.01; ***p < 0.005.

## 4. Results

### 4.1. 3D PEC Scaffold characterization and seeding solution efficiency

The process of developing the *in vitro* alginate-PEC (A-PEC) scaffold model aimed to encapsulate circulating cells within a three-dimensional (3D) scaffold, simulating the stiffness of the microenvironment of tissues affected by liquid tumors, such as chronic lymphocytic leukemia (CLL). Figure 1 illustrates the principle of this experimental setup: sterile-filtered alginate solution containing patient cells was seeded on the PEC-scaffold structure, followed by centrifugation at 400 g for 1 min. Calcium ions inherent to the PEC scaffold enabled the *in-situ* gelation of the added alginate solution, effectively encapsulating the cells within the A-PEC scaffold. This gelation process was critical to maintain the cells in a stable system.

**Figure 1.**
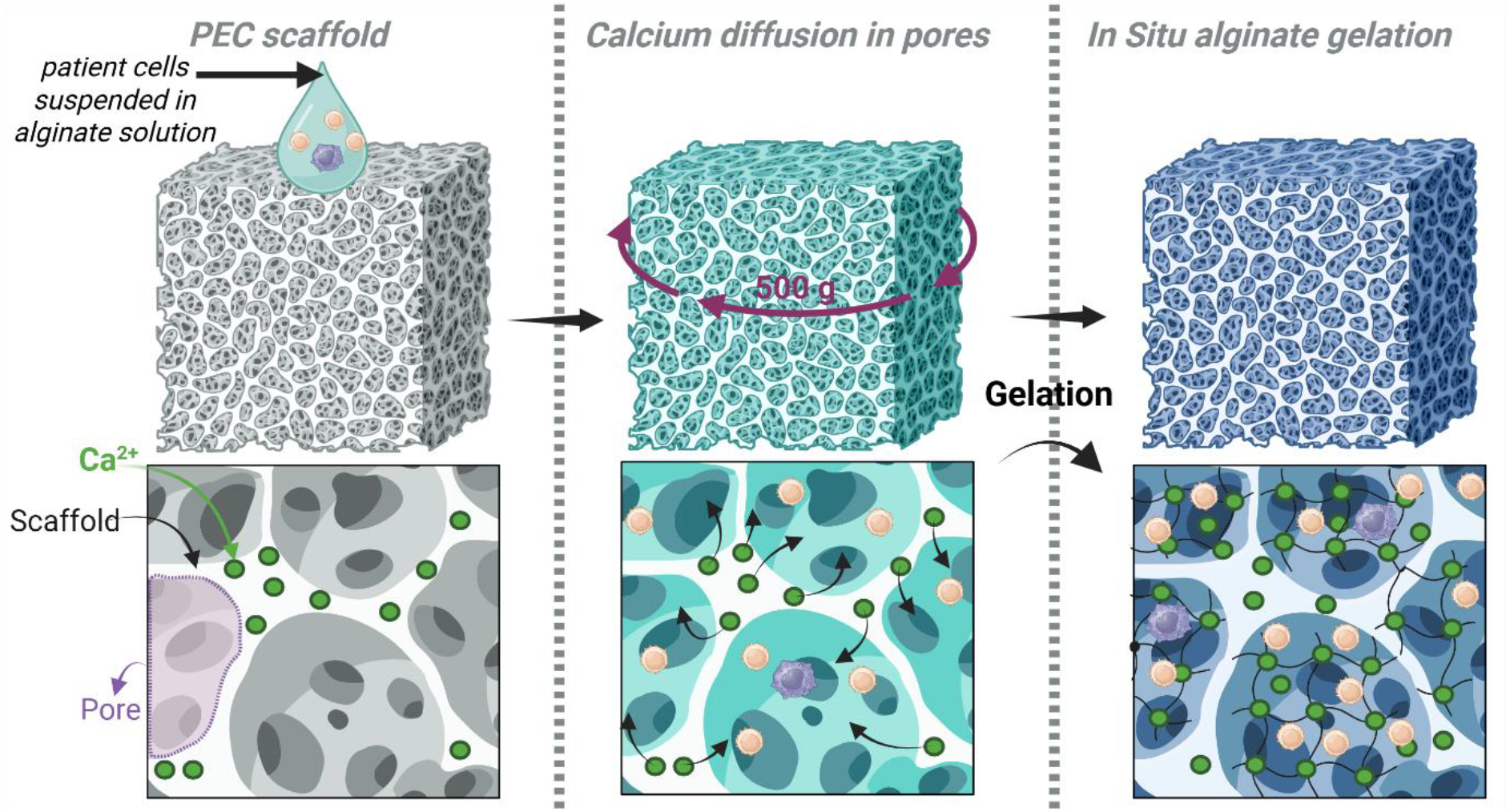
Schematic of the developed in-vitro model and its seeding method. a: Macroporous alginate-chitosan polyelectrolyte complex (PEC-) scaffold seeded with cells suspended in dilute low viscosity alginate (A) solution; b: Calcium ions diffusion from the PEC-scaffold network to the A seeding solution inside the macropores; c: In situ-gelation of A seeding solution entraps cells within the scaffold 3D structure, leading to gelled low viscosity alginate/ PEC (A-PEC) scaffold.

Figure 2 brings together the results of the characterization methods aimed at studying the properties of the scaffold and the seeding solution, and at evaluating the diffusion and gelation capacities of the LVA solution inside the pores of the scaffold. Figure 2-a shows X-ray microtomography results of the dry PEC-scaffold, revealing its overall structure. The two cross-sectional views at different depths confirm the uniform porosity throughout the entire sample. This uniformity is crucial, as it suggests that the homogeneous porosity of the PEC structure allows for a consistent diffusion of the LVA solution throughout the scaffold.

**Figure 2.**
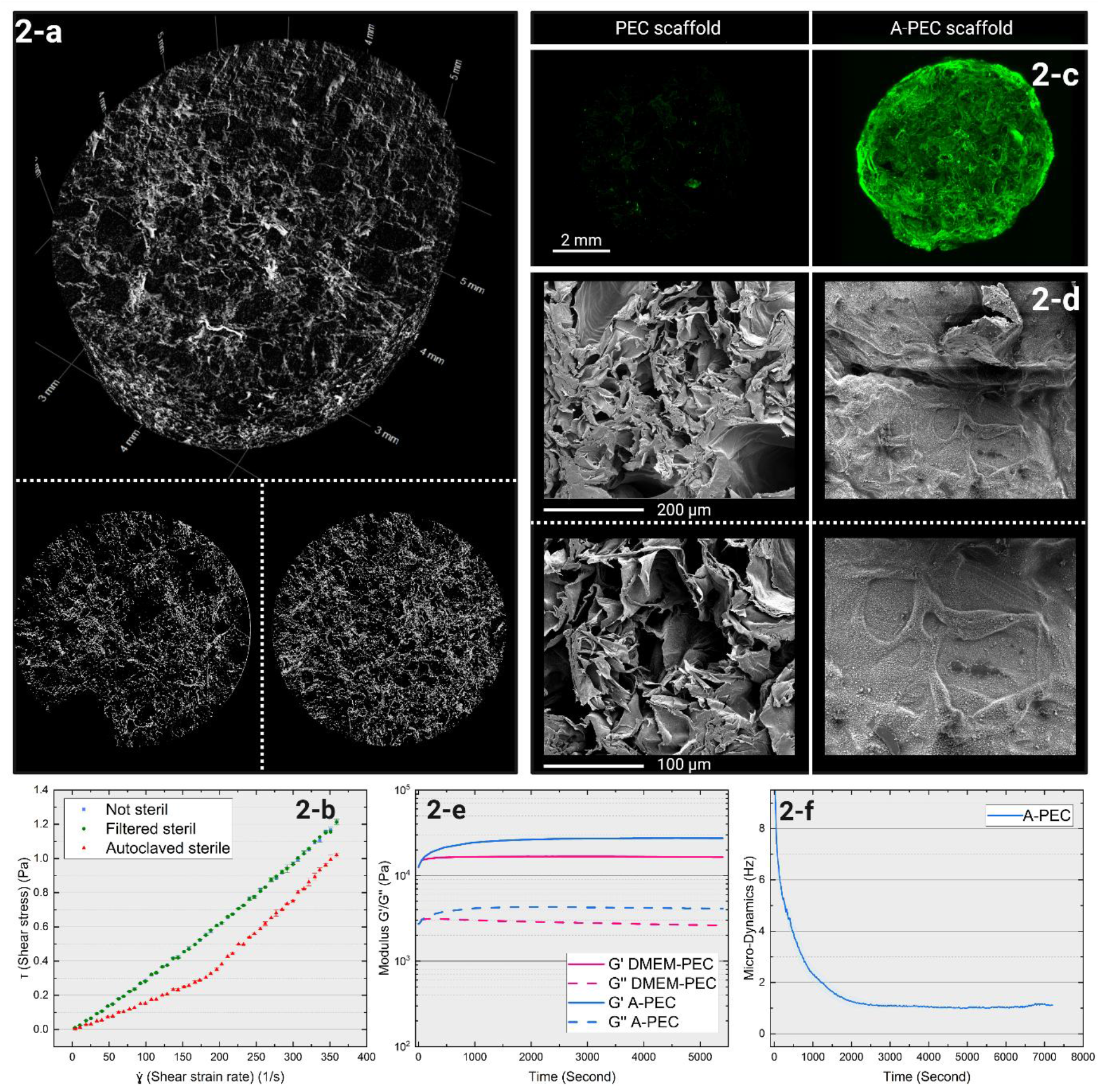
Low-viscosity Alginate (LVA) seeding solution rheological properties, ability to diffuse into PEC-scaffold macropores and in-situ gelation. 2-a: X-ray tomography of the PEC-scaffold, showing its macroporous structure before seeding with LVA solution; 2-b: Impact of 0,22 µm filtration, and autoclave sterilization on the viscosity of A seeding solution; 2-c: Confocal microscopy of PEC and A-PEC scaffold: evidence of effective diffusion of sterile FITC-labelled LVA seeding solution within the pores of the PEC scaffold; 2-d SEM images of PEC- and A-PEC scaffolds, showing pore occlusion in A-PEC scaffold, due to LVA seeding solution gelation inside the pores; 2-e: Rheological time sweep monitoring of the in situ gelation of LVA solution within the PEC-scaffold, comparing the storage (G’) and loss modulus (G’’) of PEC and A-PEC scaffolds; 2-f: Microrheology monitoring of the in situ gelation of LVA seeding solution in the PEC scaffold pores.

To develop the seeding conditions, our first goal was to investigate two key factors: (i) identifying a sterilization process that ensures a 0.5% LVA solution remains sterile without significantly altering its viscosity; (ii) assessing the gelation dynamics of the LVA solution when integrated into the PEC scaffold, through calcium ions diffusion from the scaffold matrix into the low-viscosity alginate to promote gelation.

Rheological properties of sterilized 0.5% w/w LVA solutions were assessed. The data (Figure 2-b) reveal that the viscosity of the alginate solution remained unchanged when sterilized using a 0.22 µm filter, exhibiting an identical viscosity curve to the non-sterilized control. However, autoclaving significantly reduced the viscosity of the alginate solution, suggesting mechanical degradation. This indicates that filtration is a more appropriate sterilization method for low-viscosity alginate solutions, as it preserves the solution’s mechanical properties.

To further demonstrate this penetration, Figure 2-c shows the successful incorporation of sterile low-viscosity alginate marked with FITC fluorescent dye into the scaffold. The negative control, which was devoid of alginate, exhibited negligible autofluorescence. In contrast, the scaffold loaded with FITC-marked alginate displayed a homogeneous distribution of fluorescence across the entire structure, indicating effective integration of the alginate at a macroscale level.

Alginate diffusion into the PEC pores was also quantified to assess the quality of the alginate’s incorporation at the individual pore scale. In Figure 2-d, the negative control (PEC without alginate) clearly showed empty pores, allowing us to visualize their shape, while the scaffold containing alginate demonstrated significant pore filling. This was confirmed at two different magnifications, showing the extent of alginate diffusion. These findings collectively highlight the successful and homogeneous incorporation of the LVA solution into the PEC’s pores, both at the micro- and nanostructural levels.

Next, we evaluate the mechanical properties of PEC scaffolds infused with 0.5% LVA, specifically looking at the storage modulus (G’) and loss modulus (G”). For comparison, PEC scaffolds hydrated with media alone served as a control. The data (Figure 2-e) show that the scaffolds containing gelled alginate had a higher storage modulus than the control, indicating good *in situ* gelation of the alginate within the scaffold. This increase in mechanical strength suggests a robust gelation process following the diffusion of calcium ions from the scaffold matrix into the alginate.

Finally, we used microrheology measurements to track the molecular dynamics of the alginate solution after its incorporation into the PEC scaffold. This approach allowed us to monitor the gelation process in real-time. The data (Figure 2-f) show that the molecular agitation, initially high, progressively settled over a period of 30 minutes, suggesting that gelation of the alginate solution was complete within this timeframe. This provides an estimate for the gelation time, indicating that the alginate undergoes efficient cross-linking with calcium ions within 30 minutes of being introduced into the PEC scaffold.

These findings collectively demonstrate that a 0.22 µm filtration method is ideal for sterilizing 0.5% LVA solutions without compromising viscosity, and that the *in situ* gelation process occurs effectively within 30 minutes when incorporated into the PEC scaffold.

### 4.2. Cell Seeding and Encapsulation Efficiency in the A-PEC Scaffold Model

Then we investigate two main aspects: the distribution of cells within the PEC matrix according to the presence of low viscosity alginate in the seeding solution, and the biocompatibility of the encapsulated cells over time (Figure 3).

**Figure 3.**
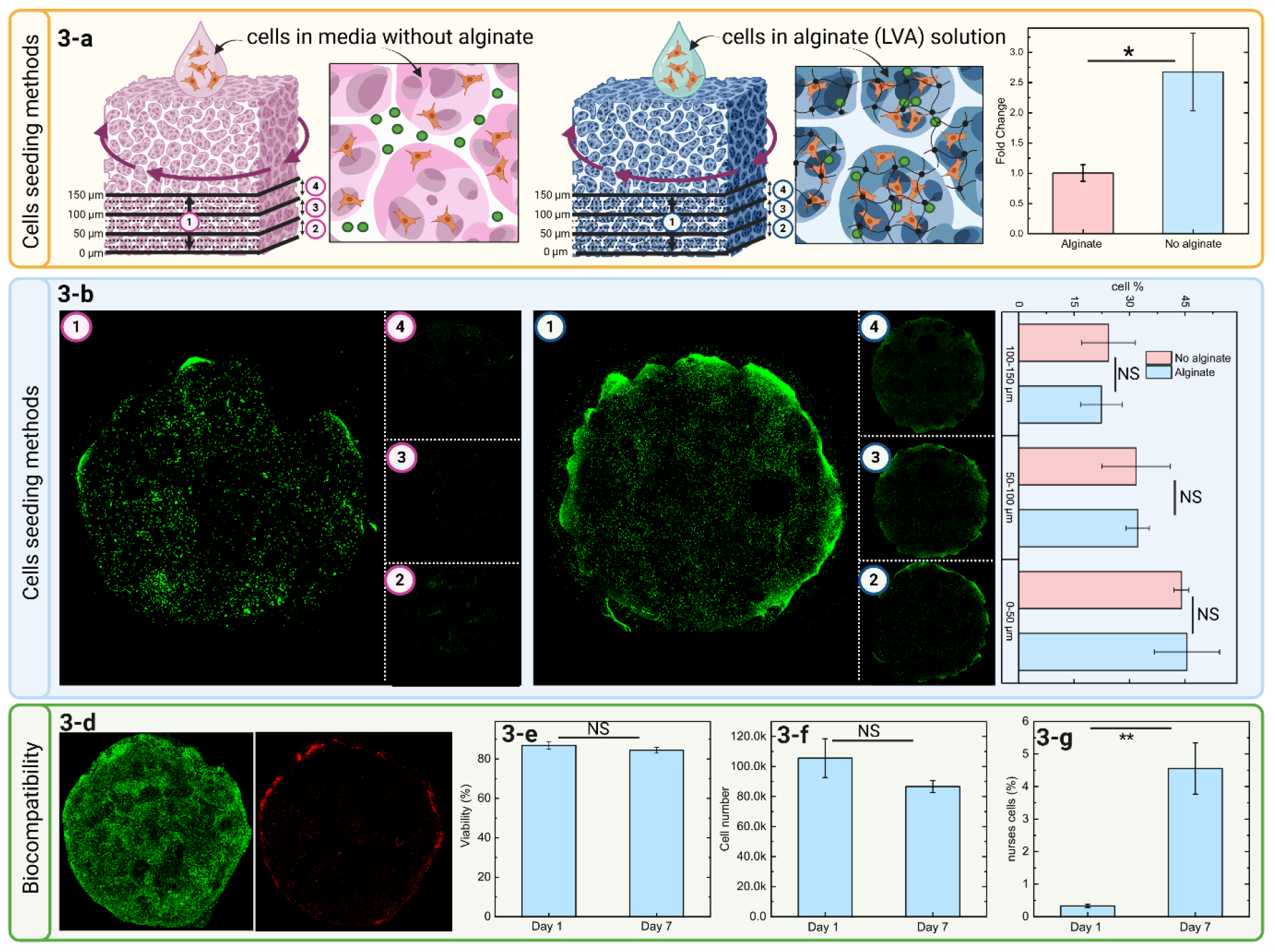
Comparison of cell seeding methods in PEC-scaffolds without and with alginate (A) in the seeding solution, and biocompatibility study in A seeded PEC scaffolds (A-PEC scaffolds) 3-a: Schematic of the cell seeding method in scaffolds, without (PEC scaffold) and with alginate (A-PEC scaffold), alongside calcein-AM staining of cells in both conditions; Cell count comparison between PEC and A-PEC scaffold, demonstrating higher cell retention in A-PEC scaffolds; 3-b: A 150 µm Z-stack confocal image (left-1), with three sequential slices (50 µm intervals) from the stack (2,3,4) shows cell distribution within the scaffold, with the quantification of cell distribution across scaffold depth, showing the percentage of cells residing in three layers: 0-50 µm, 50-100 µm, and 100-150 µm; 3-d and 3-e: Live/dead cell staining via confocal microscopy, highlighting qualitative (d) and quantitative (e) cell viability in A-PEC scaffolds (at day 1 and day 7); 3-f : Quantification of cell number from confocal images and total cell counts, demonstrating biocompatibility (at day 1 and day 7) of A-PEC scaffold. 3-g Differentiation of nurse cell populations within A-PEC scaffold model, showing stability of the nurse cell percentage across culture conditions (at day 1 and day 7).

In Figure 3a, cell quantification was achieved using microscopy across samples in defined regions of interest. The results indicate a significant increase in cell number compared to conditions without alginate, showing a 2.64-fold enhancement in cell density. This finding clearly demonstrates that our method improves the retention of circulating cells from patients with chronic cancer.

Figure 3-b demonstrates the spatial distribution of cells in the PEC scaffold under two conditions: cells suspended in culture media without alginate and cells suspended in an alginate solution. In both conditions, a centrifugation step at 400 g for 1 minute was applied to ensure proper infiltration of the liquid solutions into the PEC matrix. A full Z-stack image over a depth of 150 µm is shown (Figure 3-b1), providing a view of the cell distribution in the XY plane across the PEC scaffold. This image gives an overview of the uniformity of cell placement on the surface of the scaffold. Then, in Figures 2-a2, 2-a3, and 2-a4, the Z-stack is divided into three sections, each representing 50 µm of depth, to assess the extent of cell penetration into the scaffold along the Z-axis. This division helps to understand how the cells are distributed through different layers of the scaffold. From this analysis, we observe that the cells are well distributed across the XY plane (as seen in the full Z-stack), but the density of cells decreases progressively as we move deeper into the scaffold, which is indicated by the segmented Z-stack images (figures 3-a2 to 3-a4). Figure 3-c provides a quantification of the Z distribution observed in figure 3-a for the alginate and non-alginate conditions. Despite the alginate solution being thicker and more viscous, the data show no significant difference in cell penetration, indicating that the properties of the alginate do not adversely affect cell distribution within the scaffold.

The second part of this figure focuses on biocompatibility. Cells were seeded as described above and left 7 for days at 37°C 5% CO_2_ to ensure Nurse Like Cells (NLC) differentiation [11, 20]. Figure 3-d illustrates the excellent biocompatibility of the cells over a period of 7 days, with cell viability consistently above 80%. In figure 3-e, it is shown that while the cells do not proliferate within the matrix, they remain healthy. Finally, figure 3-f shows the percentage of NLCs, highlighting the ability of our in vitro model to support complex co-culture conditions.

### 4.3. Qualitative evaluation of Therapeutic Treatments on Cellular Viability

Our aim was to evaluate whether this new model could be used to develop drug screening with primary circulating cells from patients. In this way, we seeded fresh CLL cells from untreated patients, followed by in vitro treatments with conventionally targeted therapies. Confocal analyses using Live/dead staining permitted a qualitative assessment of cells distribution and viability within the scaffold (Figure 4).

**Figure 4.**
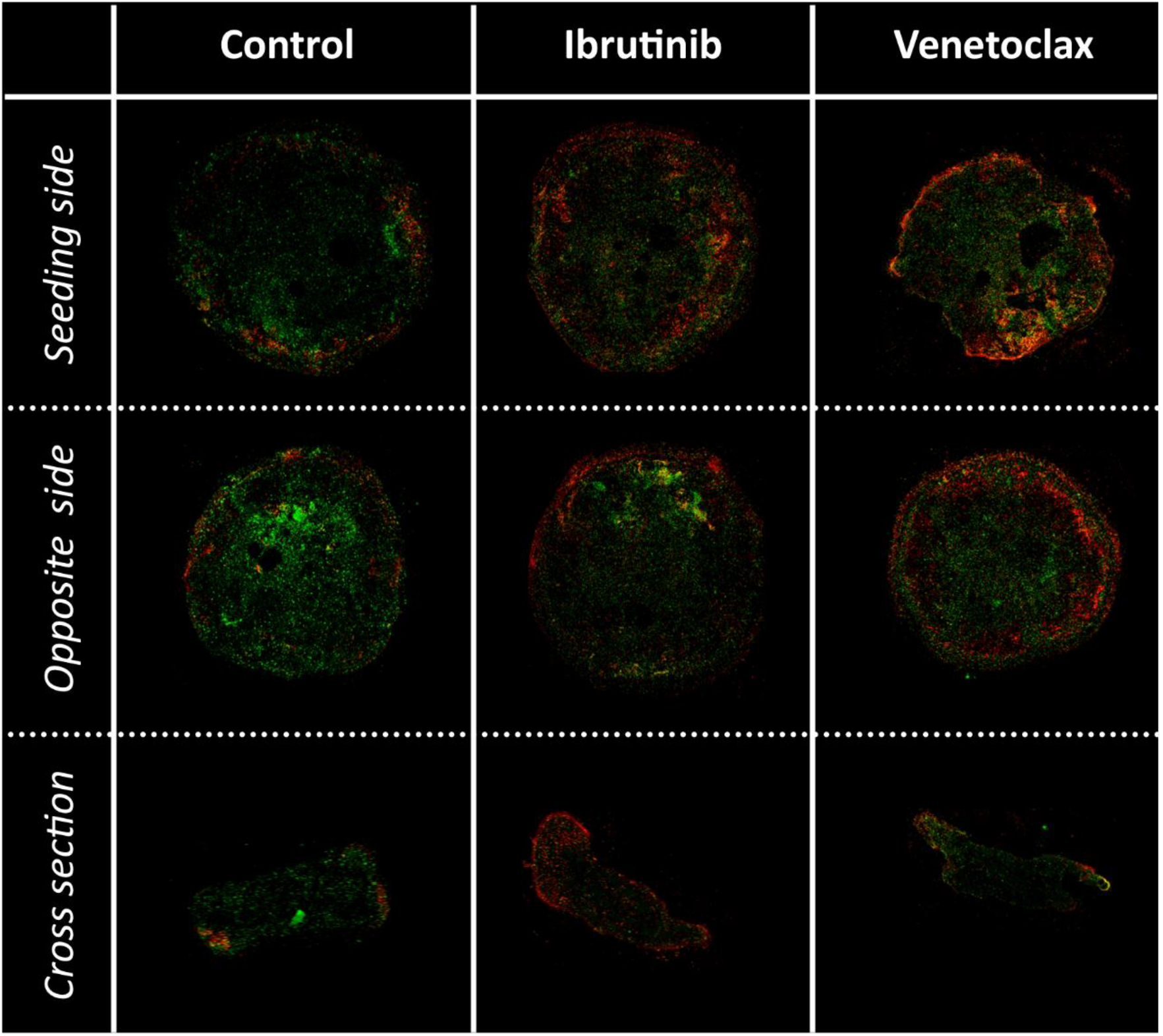
Qualitative Evaluation of CLL Cell Viability in A-PEC Scaffolds with Targeted Therapies. Confocal microscopy images depicting live/dead staining of CLL cells seeded in A-PEC scaffolds under different treatment conditions. The control (A) shows CLL cells without treatment, seeded in the A-PEC scaffold. Condition (B) presents CLL cells treated with Ibrutinib, and condition (C) shows CLL cells treated with Venetoclax, both seeded in A-PEC scaffolds. Live cells are stained green (Calcein-AM), while dead cells are stained red (EthD-1), indicating cell viability in response to treatment. The images reveal differential effects of these commonly used CLL therapies on cell survival within the 3D A-PEC scaffold environment.

Figure 4 shows live (green) and dead (red) cell staining in the PEC scaffold model for CLL, after treatment with high concentration of Ibrutinib and Venetoclax over two days. The entire scaffold was submerged in a drug-containing culture medium, ensuring equal exposure on all sides.

In the untreated control condition, cells remained predominantly viable across both the seeding and opposite sides, with minimal cell death. In contrast, following drug treatment with Ibrutinib or Venetoclax, a clear gradient of cell death emerged, with a higher concentration of dead cells on the edges. This pattern, visible in cross-section, highlighted an efficient diffusion-driven gradient of drug penetration and captured spatial patterns of drug activity on patient cells. Confocal observation of cells allowed a qualitative assessment of drug treatment efficacy although this approach may be too time-consuming and expensive for routine drug screening.

### 4.4. Quantitative assessment of Therapeutic Treatments efficacy

We then, evaluated the use of the A-PEC scaffold as a tool for drug screening in non-adherent cellular populations. We investigated the viability of chronic lymphocytic leukemia (CLL) cells under drug treatments, in the A-PEC scaffold compared with classical assays routinely used in the lab (2D culture and 3D spheroids) (Figure 5). CLL cells, obtained from blood samples of untreated patients are non-adherent and non-proliferative cells, dependent on high-density culture for their survival.

**Figure 5.**
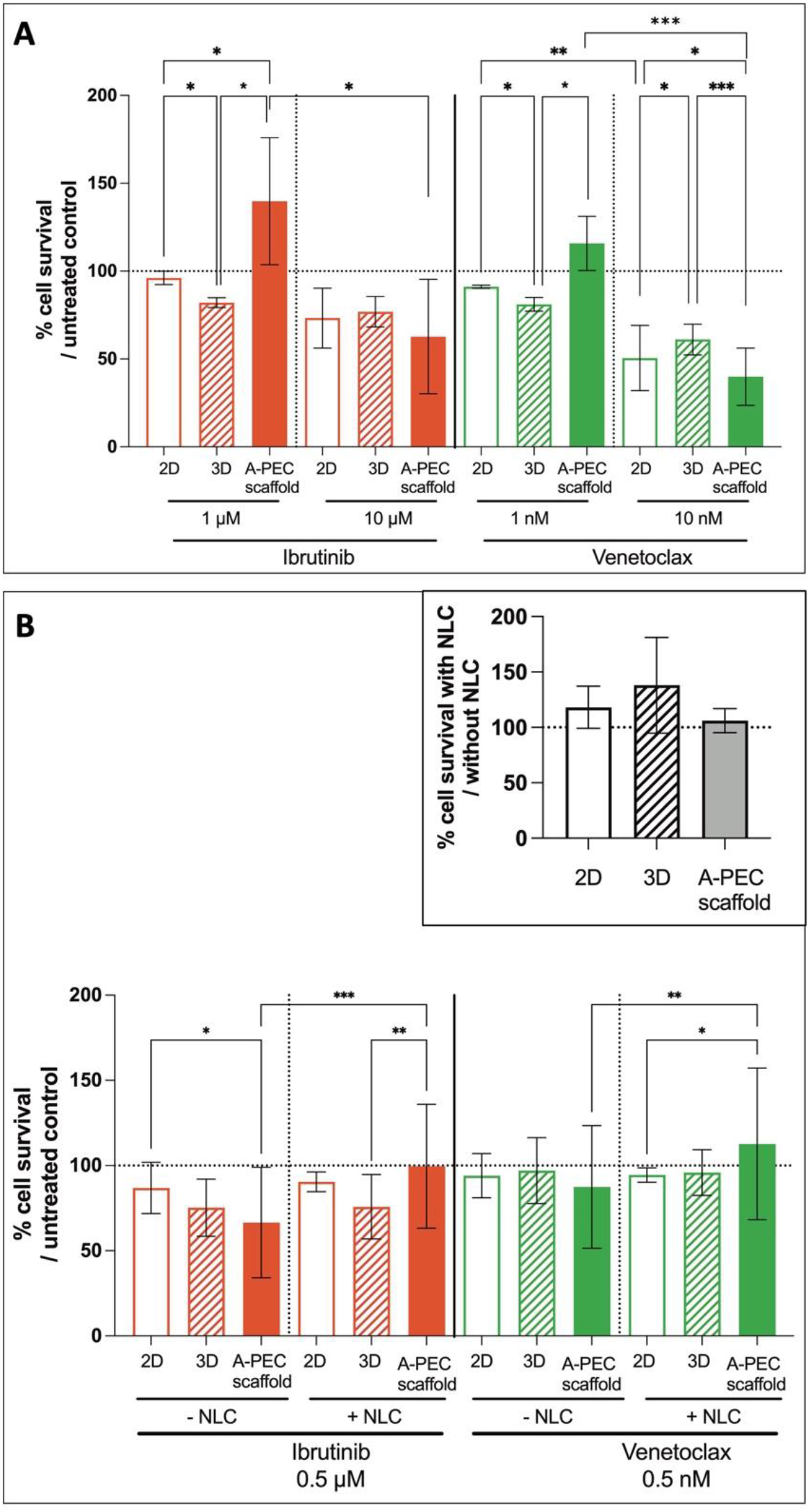
Quantification of CLL Cell Response to Ibrutinib and Venetoclax in 2D, 3D, and A-PEC scaffold culture conditions. (A) Cell viability was assessed on day 2 of culture after treatment with two concentrations of ibrutinib (1 µM and 10 µM) and venetoclax (1 nM and 10 nM). (n=4) (B) Cell viability was quantified following 7 days treatment with ibrutinib (0.5 µM) and venetoclax (0.5 nM) under the same three culture conditions in the presence or not of NLC. (n=4). Insert: % of cell viability in control culture with NLC (compared to cultures without NLC). For all experiments, % of viable cells was quantified based on negative and positive controls across the three conditions as described in Material and Methods (2D culture, 3D spheroid culture, A-PEC scaffold).

First, experiments were conducted to evaluate the dose-effect of two drugs widely used in clinical practice (ibrutinib and venetoclax at low and high doses) in a short-term culture (48 h). As observed in Figure 5-a, all culture conditions could be used to quantify a dose effect of therapeutic drugs. Indeed, both inhibitors induced a decrease in cell survival at high dose independently of culture conditions. Moreover, at low dose, results showed a better survival in the A-PEC scaffold compared with the other configurations, which can be explained by the structural complexity of the A-PEC scaffold leading to hampered drug diffusion. This assumption is consistent with what is described in CLL lymph node patients, in which cell-related density influences drug diffusion and efficacy [10].

CLL leukemic cells are highly dependent on the microenvironment, such as in lymph nodes in which cellular contact with nurse-Like Cells (NLC) ensures their better survival. Indeed, long-term culture at high density of fresh PBMCs from CLL patients induced the differentiation of NLCs, supporting B leukemic cell survival. We then assessed whether this mechanism also occurs in the A-PEC scaffold. At first, long-term culture allowed the differentiation of NLCs (Figure 3-f) in the A-PEC scaffold and the presence of NLCs ensured a good cell viability after 2 weeks (Figure 5-b insert). Then, cells were exposed to low doses of ibrutinib or venetoclax for 7 days in the presence or absence of NLCs in the different culture conditions. As shown in Fig. 5-b, NLCs protected CLL cells against both inhibitors, notably in the A-PEC scaffold.

All these results constitute proof-of concept that the A-PEC scaffold can mimic, at least in part, the cellularity of CLL lymph nodes and can be used, even in long-term cultures, as a tool to evaluate drug efficacy in the context of hematological malignancies, in parallel with 2D cultures mimicking the blood compartment.

## 5. Discussion

Although inexpensive and rapid, 2D cell culture is unable to mimic the *in vivo* 3D tumor tissue microenvironment and often fails to predict the most effective therapeutic strategy for patient treatment. The development of 3D models that yield better results is an active field of research, with scaffold-free and scaffold-based approaches used to better integrate the tumor microenvironment and the influence of stiffness on cancer cell responses and behavior [3, 4, 22] . However, applying such 3D approaches to haematological cancers is particularly challenging due to the small size, non-adherence, and potentially non-proliferative properties of patient cells.

In this context, this study focuses on the development of a 3D culture model dedicated to drug screening for liquid cancers. The challenge addressed was to effectively retain small, non-adherent cells within a 3D scaffold with tailorable mechanical properties, without affecting cell viability, and to propose a fast and effective tool for drug screening. To that aim, we developed a sterile seeding solution that permits trapping circulating tumor cells within a macroporous 3D scaffold, and we provided a proof of concept in the context of chronic lymphocytic leukemia (CLL).

The alginate biopolymer, with its ability to gel in the presence of calcium or other divalent ions, appears to be an ideal candidate for this purpose. *In situ* gelation of an alginate solution has already been proposed to trap cells within alginate-based scaffolds pores [23, 24] . However, alginate hydrogels are relatively fragile and difficult to handle. The process of gelling their pores by relying on calcium ion diffusion out of the polymer network can further weaken them.

Combining alginate and chitosan in the form of polyelectrolyte complexes can produce scaffolds with optimized and adaptable mechanical properties, particularly by adjusting the operating conditions used to produce these scaffolds. The intrinsic characteristics of the biopolymers used, such as their molecular weight (Mw), as well as the proportions between the two polysaccharides, are parameters that can modulate their mechanical properties and cellular responses [15, 25-27]. Furthermore, the drying techniques and conditions used directly affect the porosity of the scaffolds and its 3D distribution, which offers multiple possibilities for adapting to cellular needs. Some authors use freeze-drying [6, 16, 28, 29]; we exploit supercritical CO_2_ drying, which allows better preservation of nanoporosity and therefore the diffusion capacities of nutrients and waste [30]. Various porous architectures can be obtained through the use of porogens and foam formation Nevertheless, these promising structures do not allow the retention of small non-adherent cells and thus require the development of suitable seeding solutions. Because alginate and chitosan are combined in the form of a particularly stable polyelectrolyte complex, it is possible to use an alginate solution as a seeding medium and to consider gelling this solution via calcium ions included in the scaffold without overly weakening it. However, it is not obvious that calcium ions in the scaffold are present and available in sufficient quantities to allow gelling. Indeed, the research group of Zhang, which developed alginate and chitosan scaffolds through freeze-drying (without foaming or supercritical CO_2_ drying), was only able to gel alginate in the pores of its scaffolds by adding extra calcium ions [31]. Aside from the fact that this addition represents an extra step in the implementation process, it is not ruled out that the addition of calcium to the scaffold may induce the formation of a gelled layer around the scaffold, without guaranteeing gelling in the core of its structure. Under these conditions, the addition of calcium may result in the formation of a gel that resembles a sustained-release matrix and may slow down fluid circulation, not to mention that a calcium excess can affect cell viability and fate [32]. In light of the literature, it is therefore not obvious that the use of low-viscosity alginate without adding any other step or additive can allow gelation of the pores and the immobilization of cells without altering their viability or their ability to form nurse cells.

Our hypothesis was that a diluted alginate solution, containing a high cell concentration, could homogeneously diffuse and then gradually gel within the pores of a polyelectrolyte complex (PEC) scaffold without the use of additives, and could then serve as an *in vitro* liquid-cancer drug-screening platform.

Therefore, we initially focused on developing a seeding solution that could be easily sterilized and had sufficiently low viscosity to allow infiltration of the 3D network by cells through simple centrifugation. To visualize this phenomenon, we labelled the LVA with FITC and observed its localization in the pores using confocal microscopy. The observations confirm the filling of the pores, as do the SEM images of the surface and cross-section of our scaffolds. Subsequently, we monitored the gelation of the LVA solution using two techniques. Rheology measurements, following a protocol similar to that of Andersen et al., show an increase in the elastic modulus, which indicates a rapid improvement in the mechanical properties of the scaffold due to the LVA gel formation in its pores [23, 33]. Microrheology measurements have been used to confirm these results and even quantify the gelation time to around 20 minutes.

At the cellular level, the results show a distribution of cells deep within the matrix, maintenance of patient cell viability over 7 days, and the retention of these cells ‘ability to transform into nurse cells. These two factors are particularly important as they may impact the pre-treatment of cells for drug screening. The gelling of the pores of the PEC scaffold by the addition of low-viscosity alginate, without the addition of additives and without an additional step, thus allows effective cell retention without affecting their viability and main functionality. Furthermore, this gelation does not hinder confocal imaging, which allows the observation of cells and a qualitative assessment of the impact of a treatment on the distribution between dead and live cells, through Live/Dead staining.

However, in the context of treatment screening, clinical expectations are rapid responses, within 24 hours, in order to make decisions on therapeutic strategies to be undertaken for patients with recurrent CLL. We thus tested the use of an Alamar Blue kit after 24 hours of culture to see whether this tool could allow a rapid readout of the effectiveness of an immunological treatment on non-proliferating cells from patients with CLL. The results we obtained clearly demonstrate the relevance of our matrices, whose responses are different from those obtained in 2D and in 3D spheroids developed at the Cancéropole in the Toucan 2 unit. These results highlight the importance of developing 3D models that can mimic lymph nodes, complementing 2D models that more closely resemble the behavior of circulating cells in blood vessels.

The model we propose is particularly interesting. It addresses the limitations of conventional 3D hydrogel systems, in which a single material must simultaneously provide mechanical strength and cell immobilization, often resulting in networks that hinder cell–cell and cell– microenvironment communication. This approach instead combines the advantages of both systems: a porous scaffold provides mechanical support and structural organization, while a weakly crosslinked hydrogel enables cell immobilization without restricting intercellular interactions. The use of a kit like Alamar Blue allows a rapid quantitative assessment of the effectiveness of treatments on patient cells. Since the scaffold is initially presented in a dry form, it can be easily stored and used on demand; its seeding does not require any additional steps. As reported for similar scaffolds based on polyelectrolyte complexes, it can be produced in large quantities as it is obtained by casting, making it easy to present in culture plates with different well sizes, allowing rapid seeding and batch testing [6]. It can also be adapted, again by casting, to a reactor allowing dynamic-mode culture [5]. Recently, models obtained through bioprinting are multiplying [34, 35]. Compared to these models, these matrices represent a lower cost and on-demand use, which does not require waiting for the printing of a wet product with a limited shelf life.

The scaffold retains residual calcium ions, which enable in situ gelation of alginate when introduced into the scaffold. This in situ cross-linking encapsulates tumor cells in a stable 3D matrix, improving cell retention while providing a mechanically stable environment for cell survival. The PEC structure allows a large range of tunable stiffness, enabling the model to mimic the diverse stiffness characteristics seen in various soft tissues relevant to liquid tumors. This tunability positions the scaffold as a promising platform for studying the diverse behaviors of liquid tumors across various organ contexts, supporting personalized medicine applications.

Future work should include validation on larger patient cohorts, with cross-comparison of scaffold responses to known in vivo patient outcomes, to strengthen translational relevance and consolidate the model as a valuable complement to 2D systems that mimic the blood compartment.

To conclude, this 3D PEC scaffold and its dedicated seeding solution present significant advantages to be used as *in vitro* model for liquid tumors. Its tunable mechanical properties, biocompatibility, and ability to support complex cell cultures make it a powerful tool for both preclinical research and therapeutic testing. Furthermore, its ease of use and adaptability to large-scale screening enable efficient evaluation of patient-specific responses to various treatments, positioning it as a relevant platform for future therapeutic and clinical applications.

## Acknowledgements

Eloise GIMBERT for ALAMAR Blue experimentations, CMEAB for SEM imaging, Christophe Tenailleau and Benjamin Duployer for microtomography acquisition, Roland Ramsch from Speclz (formerly Formulaction) for RMaster measurements.

## Declaration of AI and AI-assisted technologies in the writing process

Statement: During the preparation of this work the author(s) used Notebook LM in order to improve readability and language. After using this tool/service, the author(s) reviewed and edited the content as needed and take(s) full responsibility for the content of the publication.

## Notes

### Competing Interest Statement

The authors have declared no competing interest.

